# The gut microbiome influences host diet selection behavior

**DOI:** 10.1101/2020.07.02.184382

**Authors:** Brian K. Trevelline, Kevin D. Kohl

## Abstract

Diet selection is a fundamental aspect of animal behavior with numerous ecological and evolutionary implications. While the underlying mechanisms are complex, the availability of essential dietary nutrients can strongly influence diet selection behavior. The gut microbiome has been shown to metabolize many of these same nutrients, leading to the untested hypothesis that intestinal microbiota may influence diet selection. Here we show that germ-free mice colonized by gut microbiota from three rodent species with distinct foraging strategies differentially selected diets that varied in macronutrient composition. Specifically, we found that herbivore-conventionalized mice voluntarily selected a higher protein:carbohydrate ratio diet, while omnivore- and carnivore-conventionalized mice selected a lower P:C ratio diet. In support of the long-standing hypothesis that tryptophan – the essential amino acid precursor of serotonin – serves as a peripheral signal regulating diet selection, bacterial genes involved in tryptophan metabolism and plasma tryptophan availability prior to the selection trial were significantly correlated with subsequent voluntary carbohydrate intake. Finally, herbivore-conventionalized mice exhibited larger intestinal compartments associated with microbial fermentation, broadly reflecting the intestinal morphology of their donor species. Together, these results demonstrate that gut microbiome can influence host diet selection behavior, perhaps by mediating the availability of essential amino acids, thereby revealing a novel mechanism by which the gut microbiota can influence host foraging behavior.

**SIGNIFICANCE:** The behavior of diet choice or diet selection can have wide-reaching implications, scaling from individual animals to ecological and evolutionary processes. Previous work in this area has largely ignored the potential for intestinal microbiota to modulate these signals. This notion has been highly speculated for years but has not yet been explicitly tested. Here we show that germ-free mice colonized by differential microbiomes (from wild rodents with varying natural feeding strategies) exhibited significant differences in their voluntary dietary selection. Specifically, differences in voluntary carbohydrate selection were associated with plasma amino acid levels and bacterial genes involved in the metabolism of tryptophan. Together, these results demonstrate a role for the microbiome in host nutritional physiology and behavior.

## INTRODUCTION

Proper nutrition is essential to life, and thus animals have evolved complex internal sensory systems that help maintain nutritional homeostasis by regulating macronutrient intake^1^. The intestinal tract plays a critical role in this process by liberating dietary nutrients (*e*.*g*., essential amino acids) that communicate meal quality to the central nervous system by direct stimulation of enteric nerves or through post-absorptive peripheral signals^2–4^. The intestinal tract also harbors trillions of microorganisms (collectively known as the gut microbiome), which have been shown to influence numerous aspects of host behavior, most likely through metabolites that interact with host sensory systems^5^. Given the importance of dietary nutrients in the regulation of food intake and diet selection^6^, the gut microbiome may influence host foraging behavior through metabolic processes that affect the availability of nutrients (or their derivatives) recognized by the central nervous system^2,7,8^. For example, a recent study showed that experimental colonization of *Providencia* bacteria in the gut of the model organism *C. elegans* resulted in divergent foraging preferences through the bacterial synthesis of the neurotransmitter tyramine from the essential amino acid tyrosine^9^. While studies in model systems provide powerful opportunities to dissect host-microbe interactions^10^, the microbiome field recognizes the need to address and study the complexity of these interactions in ecologically-realistic scenarios in which animals can harbor thousands of microbial taxa^11,12^. It has been suggested that these complex microbial communities could elicit host foraging behaviors that enrich the intestinal environment in nutrients on which they depend (*i*.*e*., promoting their own fitness)^7^, while others have posited that a positive-feedback relationship between dietary nutrients and microbial community composition eventually results in stable microbial communities and host foraging behaviors^8^. However, these potential mechanisms operate under the assumption that the gut microbiome influences diet selection behavior – a hypothesis that has existed for years^7,8^, but has never been tested using complex microbial communities, or within an ecological or evolutionary context.

The transplantation of intestinal microbiota into germ-free mice is a powerful approach for disentangling the effects of the gut microbiome on host phenotypes from other potentially confounding factors (*e*.*g*., host genetics)^13^. This approach has been successfully applied using a wide range of donor species (*e*.*g*., termites, zebrafish)^14^, demonstrating that germ-free mice are a tractable model system for understanding the function of gut microbiota in evolutionarily-distant organisms. In one notable example, Sommer *et al*. used fecal microbiome transplants from brown bears into germ-free mice (two species separated by ∼94 million years of evolution) to show that seasonal changes in gut microbiota influence host energy metabolism^15^. In our study, we used this approach to determine whether the gut microbiome influences diet selection behavior. We chose three rodent species with distinct foraging strategies as microbial donors for germ-free mice: a carnivore/insectivore (southern grasshopper mouse, *Onychomys torridus*), an omnivore (white-footed mouse, *Peromyscus leucopus*), and an herbivore (montane vole, *Microtus montanus*). These three species are in the same taxonomic family (Cricetidae) and are all equally distantly related to lab mice (∼27 MYA; *Mus musculus*, family Muridae)^16^. Under sterile laboratory conditions, we randomly divided 30 adult male germ-free mice into Carn-CONV, Omni-CONV, and Herb-CONV treatment groups (n = 10 mice per group), where each mouse in a given group was “conventionalized” (*i*.*e*., inoculated) with the cecal contents of a unique, wild-caught donor individual (to better reflect natural interindividual variation) (Fig. 1a). One recipient mouse from the Herb-CONV group was excluded from our dataset due to aberrant behaviors that indicated possible injury during microbiome transplants. Conventionalized mice were acclimated to their microbiota for 7 days, during which they were offered only sterile water and a low protein:carbohydrate ratio diet (LPC; Table S1). There were no differences in daily or cumulative macronutrient and food intake across treatment groups during the acclimation period (Fig. S1; Dataset S1). After acclimation, conventionalized mice were given a choice between the LPC diet and one with a higher P:C ratio (HPC; Table S1) for a period of 11 days (Fig. 1a). Importantly, these diets had identical energy densities (caloric content per gram).

**Fig. 1.**
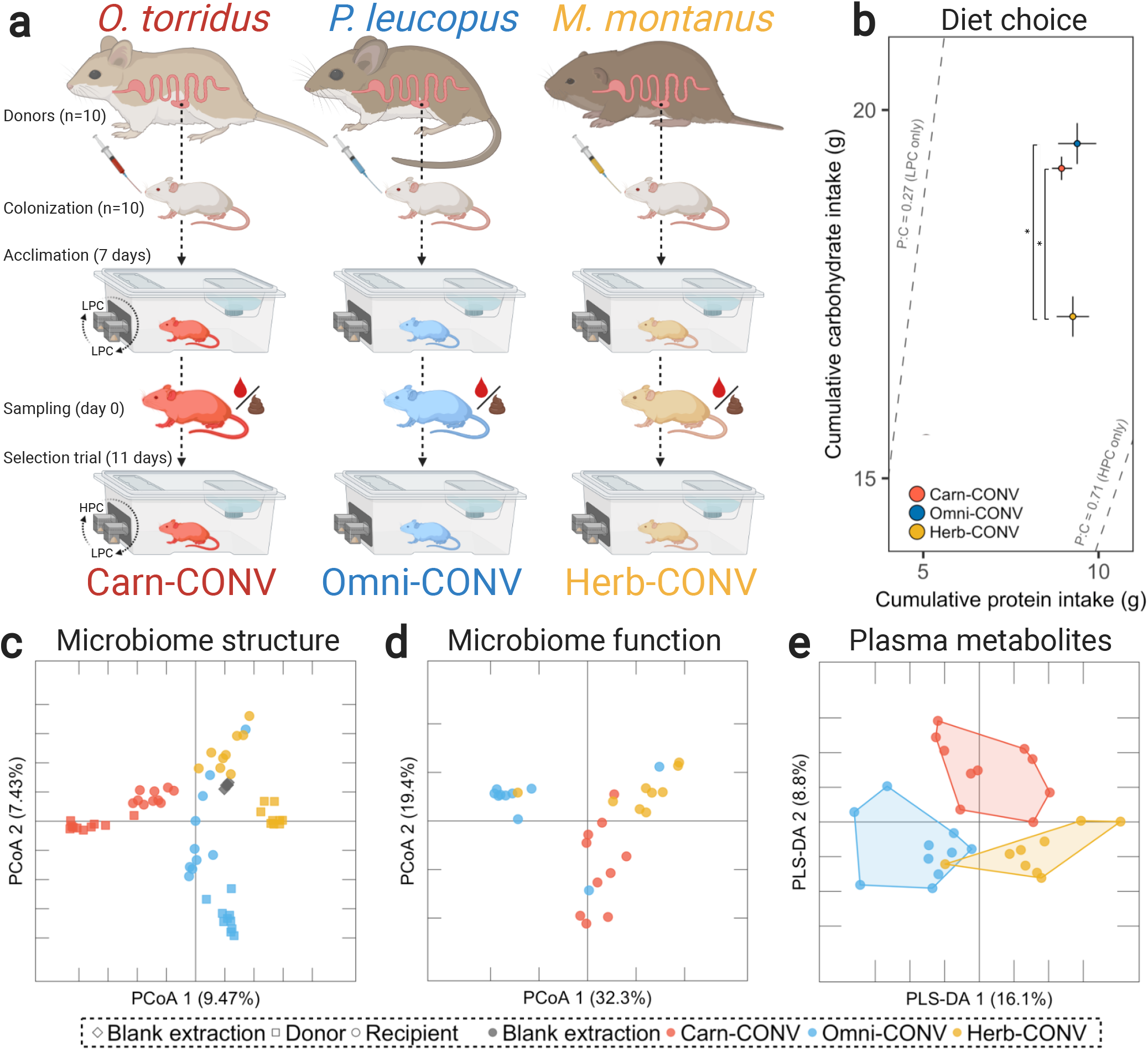
The gut microbiome influences host diet selection behavior. **a**, Overview of experimental design. Germ-free mice were colonized with the gut microbiome of three species of wild rodents with distinct foraging strategies: carnivorous *Onychomys torridus* (Carn-CONV), omnivorous *Peromyscus leucopus* (Omni-CONV), and herbivorous *Microtus montanus* (Herb-CONV). Conventionalized mice were acclimated on LPC diet for 7 days before day 0 blood and fecal sampling. After acclimation, conventionalized mice were then given a choice between LPC and HPC diets for 11 days. Daily diet intakes were tracked via two feeder hoods, which were rotated daily to avoid learned preferences. **b**, Treatment groups differed significantly in macronutrient intake (Wilks’ λ = 0.455, Cohen’s *f*^*2*^ = 0.41, power = 0.98, P = 0.0007), with Herb-CONV mice voluntarily consuming fewer carbohydrates than the Omni- and Carn-CONV groups (F = 9.22, P = 0.001). There was no difference in cumulative protein intake across treatment groups (F = 1.362, P = 0.275). Dashed rails and associated P:C ratios indicate the expected result if mice consumed only a single diet. Error bars represent the standard error of the mean (SEM). **c**, Principal coordinate analysis (PCoA) of 16S rRNA inventories of wild donors (squares) and conventionalized recipients at day 0 (circles) using Bray-Curtis dissimilarity. Microbial community structure differed significantly among wild donors (Pseudo-F = 7.41, P = 0.001) and recipients (Pseudo-F = 3.24, P = 0.001). All groups differed significantly from blank extraction controls (gray diamonds; Pseudo-F = 4.78, P = 0.001). **d**, PCoA analysis showing a statistically significant difference in the relative abundances of microbial KEGG modules using Bray-Curtis dissimilarity (Pseudo-F = 5.96, P = 0.001). **e**, PLS-DA analysis illustrating broad differences in identified plasma metabolites across conventionalized mice at day 0. * denotes P ≤ 0.05.

To determine whether treatment groups differed in foraging behavior, we employed a state-space approach known as the Geometric Framework in which foraging decisions are analyzed within a multi-dimensional nutritional space where each functionally relevant nutrient forms a single dimension^17,18^. In this study, we defined these nutritionally-explicit dimensions as protein and carbohydrate intake, thereby allowing us to measure the effect of the gut microbiome on host diet selection. Supporting the hypothesis that the gut microbiome influences diet selection behavior, this approach revealed statistically significant differences in macronutrient intake across groups of conventionalized mice (Fig. 1b). Treatment groups differed significantly in daily (Fig. S1) and cumulative carbohydrate intake (Fig. 1b) during the diet selection trial. Specifically, Herb-CONV mice voluntarily consumed fewer carbohydrates than Carn-CONV and Omni-CONV mice. This trend was most apparent after approximately 1 week of diet choice (Fig. S1), suggesting that it may take time for internal nutritional signals to stabilize^19^ and for associative learning^20^ to affect host feeding behavior. In contrast, treatment groups did not differ in either daily (Fig. S1) or cumulative protein intake (Fig. 1b). Lower cumulative carbohydrate intake among Herb-CONV mice led to their selection of a significantly higher P:C ratio diet compared to Omni-CONV and Carn-CONV mice (Fig. S2). Interestingly, we also observed a significant difference in total food intake among Herb-CONV mice compared to the other treatment groups (Fig. S1), suggesting that Herb-CONV mice’s preference for the higher P:C ratio diet may have permitted them to reduce total energy intake without affecting nutritional homeostasis (*i*.*e*., protein-leveraging)^19^. Under natural scenarios, such differences in selected P:C ratios could be accomplished by animals incorporating different levels of insects, seeds, or foliage into their diets. The ratio of macronutrients an animal consumes, rather than the total amount of any individual nutrient, has significant effects on animal physiology, life history, and reproductive fitness^21–23^. The preference of Herb-CONV mice for the HPC diet are also consistent with previous studies showing that *Microtus* voles prefer high-protein foods when available^24,25^, though a follow-up study on the foraging preferences of *M. montanus* with respect to specific dietary nutrients would more robustly support the ecological significance of our findings. More generally, these results are also consistent with the “nitrogen limitation hypothesis”, which posits that the relative scarcity of nitrogen in plant materials may drive the opportunistic consumption of higher protein foods among herbivores^26–28^. Interestingly, the hindgut microbiota of herbivorous mammals are also nitrogen-limited^29^, and so our findings offer support to the hypothesis that microbes may alter host foraging behaviors to enrich the intestinal environment in necessary nutrients^7^.

Next, we characterized day 0 (7 days post-inoculation and just prior to diet selection trial) gut microbial community structure, microbiome function, and plasma metabolites of conventionalized mice to determine how these aspects were associated with differential diet selection across treatment groups. 16S rRNA inventories confirmed that both donors and recipients harbored distinct bacterial communities that differed significantly from blank extraction controls (Fig. 1c; Fig. S3; Fig. S4). We observed significant differences in colonization efficiency across treatment groups. Specifically, microbial communities of Carn- and Omni-CONV recipients were significantly most similar to those of their donors, while Herb-CONV recipients were not significantly similar to any donor group (Fig. S4). It is expected that recipient communities would not match donors identically, as the *Mus* host physiology reshapes donor communities^30^, and our donor communities were collected from individuals in the wild, and thus our design does not account for the well-documented effects of captivity on the microbiome^31^. The comparatively lower colonization efficiency among Herb-CONV mice may have been driven by the low content of indigestible plant fibers that are primarily fermented by microbes. Even in established microbiomes, differences in the content or composition of dietary fiber can result in the extirpation of some fermentive microbes^32,33^. However, Herb-CONV mice were successfully colonized by donor microbiota in the phylum Firmicutes (classes Bacilli and Clostridia), notably those in the family Lachnospiracae, which are strict anaerobes known for their ability to transform plant fibers into volatile fatty acids in the mammalian digestive tract^34^. Additionally, microbiomes from herbivorous mammals colonize germ-free mice at a lower absolute density than microbiomes from omnivorous or carnivorous mammals^35^. More work is required to understand differential transfer of microbiomes across species, and we discuss this limitation in more detail below.

Bacterial ASV richness and phylogenetic diversity were similar across donor groups, but significantly lower in Herb-CONV mice compared to the other treatment groups (Fig. S4). In general, the bacterial communities of conventionalized mice were dominated by the phyla Bacteroidetes and Firmicutes (Fig. S4). Importantly, all recipient fecal samples tested negative for the presence of pathogenic microorganisms. Metagenomic analysis of recipient fecal samples revealed a statistically significant effect of donor species on the relative abundances of 183 (51%) KEGG functional modules (Fig. 1d; Dataset S2). These differences in microbiome community structure and function were accompanied by concomitant differences in plasma metabolites (Fig. 1e), with 27 identified metabolites (16%) differing significantly across treatment groups (Dataset S3). Together, these results demonstrate that interspecific differences in gut microbial communities across rodents with divergent foraging strategies translate to distinct microbial functions and metabolite profiles independent of host diet.

There is substantial evidence that the availability of circulating essential amino acids (EAAs) provide peripheral signals that act to regulate macronutrient intake and diet selection^4,6^. Despite consuming identical diets prior to the selection trial, treatment groups differed in circulating levels of several amino acids, with Herb-CONV mice exhibiting significantly higher amounts of the EAAs lysine, isoleucine, methionine, phenylalanine, and tryptophan (Fig. 2a). While EAAs are primarily derived from the diet, bacteria can also produce these peptides through their own metabolic processes^36^, and thus the gut microbiome may act as a source of EAAs for their hosts. In support of this hypothesis, treatment groups exhibited broad differences in the microbial synthesis and degradation of EAAs (Fig. 2b). Notably, the microbiome of Herb-CONV mice had a higher abundance of genes involved in the synthesis of aromatic amino acids (phenylalanine, tryptophan, and tyrosine) (Fig. 2b), all of which are synthesized from chorismate (product of the Shikimate pathway)^37^. The ratios of bacterial genes involved in tryptophan biosynthesis (M00023) to those involved in tryptophan degradation via the kynurenine pathway (M00038) were significantly correlated with plasma tryptophan (Fig. 2c). Given that conventionalized mice consumed identical diets prior to blood collections, these results demonstrate that bacterial metabolism can alter the availability of circulating levels of plasma EAAs, consistent with recent studies conducted in *Drosophila*^38^.

**Fig. 2.**
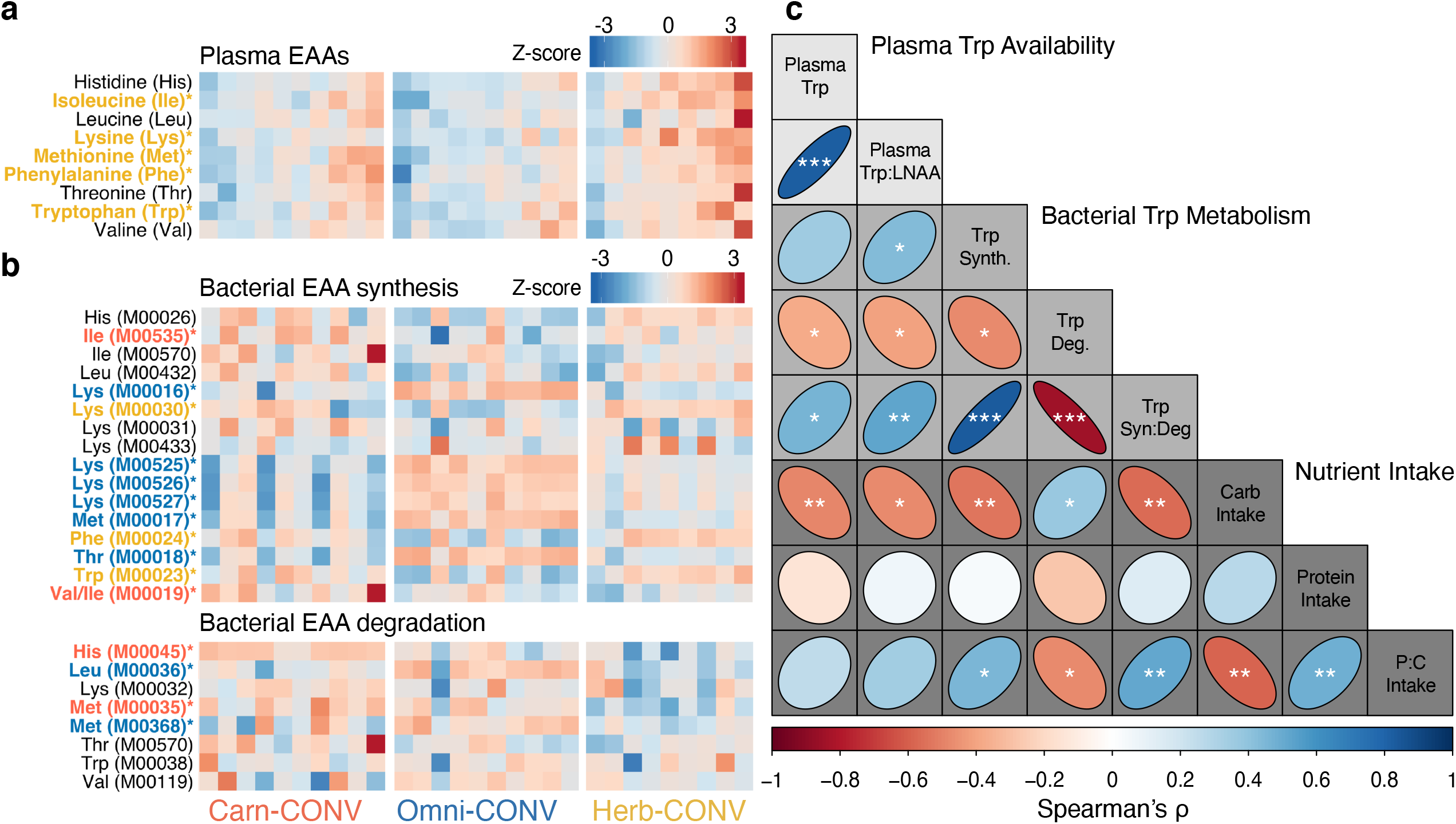
Day 0 plasma tryptophan availability and bacterial tryptophan metabolism are associated with differential macronutrient intake across treatment groups. **a**, Heatmap illustrating broad differences in plasma levels of essential amino acids across treatment groups, with Herb-CONV mice exhibiting significantly greater levels of lysine (X^2^ = 6.13, P = 0.047), isoleucine (X^2^ = 11.42, P = 0.003), methionine (X^2^ = 6.13, P = 0.047), phenylalanine (X^2^ = 6.13, P = 0.047), and tryptophan (X^2^ = 9.10, P = 0.011) compared Carn-CONV and Omni-CONV mice. Columns represent individual conventionalized mice for each treatment group. * denotes P ≤ 0.05 and color indicates the treatment group with greatest circulating plasma levels (red = Carn-CONV, blue = Omni-CONV, and yellow = Herb-CONV). **b**, Heatmap illustrating broad differences in the abundances of microbial genes associated with metabolism of essential amino acids (Dataset S2). * denotes P ≤ 0.05 and color indicates the treatment group with greatest relative abundance. **c**, Correlation plot summarizing relationships between plasma tryptophan availability, bacterial tryptophan metabolism, and host diet selection among conventionalized mice. The direction and color of the ellipses indicate whether correlations were positive or negative, and asterisks indicate whether Spearman’s correlations were statistically significant (* denotes P ≤ 0.05, ** denotes P < 0.01, and *** denotes P < 0.001).

There is emerging evidence that bacterial tryptophan metabolism is a key mechanism by which the gut microbiome can influence host behavior^39,40^. This relationship is a consequence of tryptophan’s role as the primary regulatory molecule for the synthesis of central serotonin (5-hydroxytryptamine, 5-HT)^41^, which has been shown to drive foraging behavior and diet selection in several experimental studies^42,43^. For example, when given a choice between low- or high-carbohydrate meals, rats receiving hypothalamic injections of 5-HT significantly reduced their carbohydrate intake^44^. Importantly, serotonin synthesis is extraordinarily sensitive to plasma tryptophan availability, and thus plasma tryptophan is generally considered a reliable proxy for central serotonin^45^. Therefore, we predicted that plasma tryptophan would be associated with differences in diet selection among conventionalized mice. Indeed, we found a statistically significant correlation between day 0 plasma tryptophan and subsequent voluntary carbohydrate intake (Fig. 2c). More recent work has argued that serotonin synthesis is affected by the availability of tryptophan relative to the large neutral amino acids (LNAA: Leu, Ile, Phe, Tyr, and Val) that compete for transport across the blood brain barrier^46^. Consistent with these studies, we found a statistically significant correlation between day 0 Trp:LNAA ratios and cumulative carbohydrate intake (Fig. 2c). Further, the ratio of tryptophan biosynthesis and degradation KEGG modules were also statistically significant predictors of carbohydrate and P:C intake (Fig. 2c). Overall, these results support the hypothesis that bacterial tryptophan metabolism influences host diet selection behavior.

Interspecific differences in foraging behavior are generally associated with diet-specific adaptations to intestinal physiology. For example, herbivores generally maintain an enlarged cecum (fermentation chamber) that enhances the digestibility of low-quality, carbohydrate-rich foods^47^. Given that the gut microbiome can profoundly alter host intestinal gene expression and physiology^48–50^, divergent microbial communities may drive differences in intestinal morphology across feeding strategies. At the conclusion of the diet selection trial (day 11), we quantified intestinal morphology with the prediction that conventionalized mice would exhibit differences that broadly reflected that of their donor species. While there was no change in body mass over the duration of the experiment (F = 1.01, P = 0.377), treatment groups differed significantly in empty colon mass (Fig. 3b), with Herb-CONV mice exhibiting comparatively larger colons than those in other treatment groups. There were no significant differences in cecum mass (Fig. 3a) or colon length (Fig. 3c). In general, the comparatively larger colons observed in Herb-CONV mice are consistent with evolutionary adaptations observed in herbivorous animals, which generally maintain larger hindguts to promote digestion^47^. The gut is a highly dynamic organ that can rapidly change in mass and length in response to environmental conditions, often through altered rates of cellular proliferation in intestinal crypts and cell loss through sloughing or apoptosis at the ends of intestinal villi, but also through the change of the size of individual enterocytes^51^. In the future, histological analyses could be conducted to investigate whether these changes in gut size are driven by hyperplasia (increase in cell number) and/or hypertrophy (increase in cell size), and to rule out the possibility for these differences to be driven by intestinal inflammation.

**Fig. 3.**
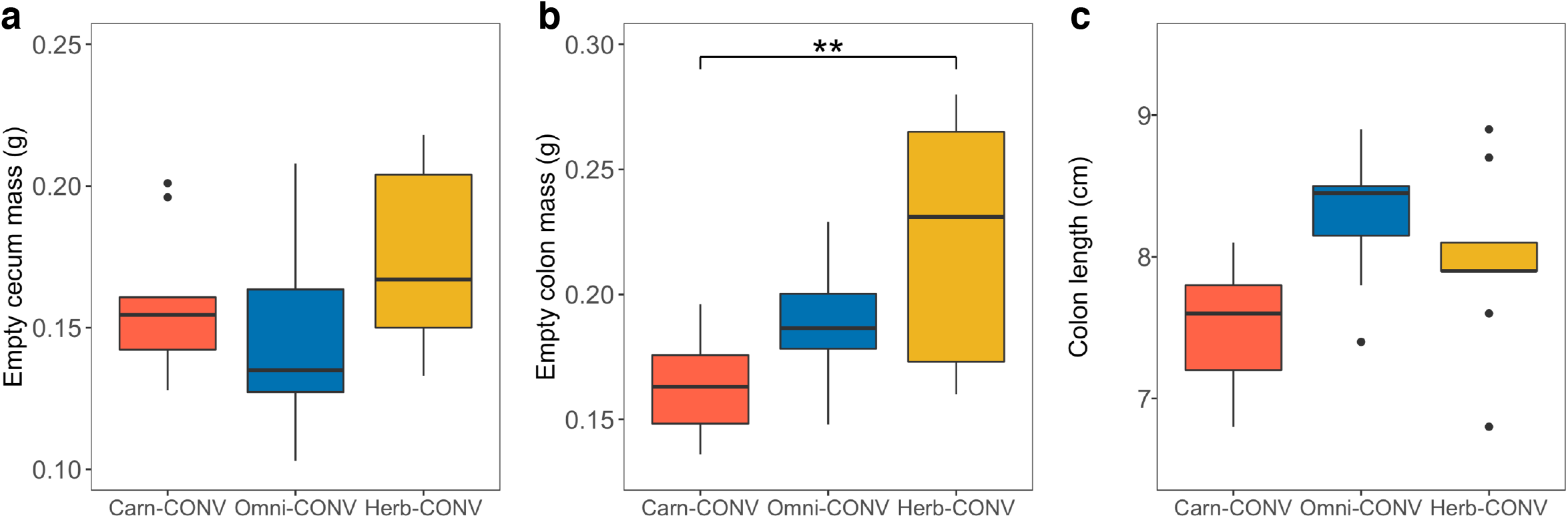
Treatment groups exhibit differences in intestinal morphology. **a**, Empty cecum mass did not differ significantly across treatment groups (F = 2.18, P = 0.133). **b**, Empty colon mass differed significantly across treatment groups (F = 6.91, P = 0.004), with Herb-CONV mice exhibiting a greater colon mass than Carn-CONV mice (FDR-adj. P = 0.003). **c**, Colon length did not differ significantly across treatment groups (F = 2.03, P = 0.151). ** denotes P ≤ 0.01.

While the observed differences in gut size are consistent with adaptations observed in herbivores, our study only tested the microbiome of a single species from each feeding strategy. A robust test of whether the microbiome recapitulates the differences in gut size observed across feeding strategies would require several donor species from each dietary strategy. Another question is whether the gut microbiome affected intestinal morphology directly or via differential diet selection. While our experimental design makes it difficult to disentangle the effects of differential diet selection from those of microbiome, it is worth noting that previous work has demonstrated that lab mice fed low P:C ratio diets had larger intestinal compartments (*e*.*g*., colon) compared to those fed higher P:C diets^50^. In our study, we observed the opposite – Herb-CONV mice, which consumed a higher P:C ratio diet (Fig. 1b), exhibited larger colon masses (Fig. 3). These results contradict the generally accepted model of adaptive physiological responses to dietary carbohydrates, suggesting that the gut microbiome may drive interspecific differences in host intestinal physiology to some extent, independent from the effects of diet and genetics.

Here, we present evidence for an effect of the gut microbiome on host diet selection behavior, however, it is important to recognize that our approach has several substantial limitations. For example, the relative differences in nutrient composition between diets have been shown to greatly influence animals’ ability to distinguish and differentially feed^19^, suggesting that our differential diet selection results may have been more pronounced if we had used diets with greater differences in macronutrient content. Further, previous work has shown that the evolutionary distance between donor species and germ-free mice can affect the efficacy of microbiome transplants^14^. While our selected donor species were similarly distant to *Mus musculus*, there were significant differences in colonization success across donor species, suggesting that cecal microbiota may be specifically adapted to their hosts. While differences in colonization efficiency may limit our ability to robustly connect our study to the ecology of donor species, this limitation should not diminish our major finding that conventionalized germ-free mice harboring compositionally and functionally distinct microbiotas differing in microbial diversity exhibited different feeding preferences. Overall, our approach is stronger than comparing conventional mice with the highly artificial state of germ-free mice, and the complex microbial communities that we used better reflect reality, which is recognized as pressing need in the field of host-microbe interactions^11,12^.

In this study, we found that conventionalized germ-free mice harboring distinct gut microbiota exhibited significant differences in diet selection behavior, providing support for our core hypothesis that microbiota can influence foraging decisions. Specifically, our study provides evidence that variation in the gut microbiota alters host nutrient availability and can yield significant differences in the diet selection of conventionalized mice in just 11 days, likely through differential bacterial metabolism and downstream availability of EAAs, especially tryptophan. These findings are largely consistent with recent mechanistic work in model systems^9,38^, but address the natural variation in microbial communities that exist among individuals and across species^11,12^. Therefore, this study not only represents a novel contribution to a large body of work showing that the gut microbiome is a key player in host physiology and performance^52^, but also more broadly supports the hypothesis that the gut microbiota can influence ecological and evolutionary processes shaping animal behavior. Foraging strategies and feeding behaviors can influence many aspects of an animal’s ecology (*e*.*g*., the need to obtaining specific nutrients while also avoiding predators^53^), and animal feeding can also shape the structures of entire plant and animal communities^54^. Thus, there may be an underexplored role for gut microbes to influence far-reaching aspects of animal and ecosystem ecology through influencing the feeding behavior of their hosts.

## Acknowledgements

This work was supported by the National Science Foundation (IOS-1942587 to K. Kohl) and a grant from the University of Pittsburgh Central Research Development Fund. We thank A. Darracq, M. D. Dearing, T. Derting, R. Martínez-Mota, and S. Secor for help with animal collections and dissections.

## Author Contributions

K.D.K conceived the project. B.K.T designed the experiments, performed the experiments, collected the data, interpreted the results, and wrote the manuscript with guidance from K.D.K.

## Additional Information

Supplementary information is available for this paper. Sequencing data are deposited in the NCBI SRA database under PRJNA629007. The authors declare no competing financial interests. Correspondence and requests for materials should be addressed to B.K.T.

## MATERIALS AND METHODS

### Wild rodents

Wild *Onychomys torridus* were collected in August 2018 from field sites in near Green Valley, Pima Co., AZ (31.802834, -110.891172), *Peromyscus leucopus* in May 2018 near Murray, Calloway Co., KY (36.686582, -88.221204), and *Microtus montanus* in July 2018 at Timpie Springs Waterfowl Management Area, Dugway, Tooele Co., UT (40.753708, -112.639903). Ten individuals from each species were collected using baited Sherman live traps under the following state permits: *O. torridus* (AZ Game and Fish Dept., SP627958), *P. leucopus* (KY Dept. of Fish and Wildlife, SC1911097), and *M. montanus* (UT Division of Wildlife Resources, 1COLL5194-2). Animals were euthanized within 12 hours and immediately dissected under IACUC protocols registered at the University of Utah (16-02011 to D. Dearing), Murray State University (2018-026 to T. Derting), and University of Alabama (18-04-1159 to S. Secor). Cecum contents for microbiome transplants were transferred to 1.7 mL Eppendorf tubes using sterile instruments and temporarily frozen at -20ºC in the field before long-term lab storage at -80ºC.

### Microbiome transplants

Donor cecum contents were diluted at 100mg/mL in sterile phosphate-buffered saline containing 0.2 g/L Na_2_S and 0.5 g/L cysteine as reducing agents^55,56^. Under sterile laboratory conditions, 30 adult (aged 6-8 weeks) male germ-free C57BL/6 mice (Taconic Biosciences, Inc., Rensselaer, NY) were randomly divided into Carn-CONV, Omni-CONV, and Herb-CONV groups (n = 10 mice per group), where each mouse in a given group was colonized by oral gavage of 200 μL of fecal slurry from a unique, wild-caught donor individual. Conventionalized mice were then singly-housed in sterile static cages (Innovive, Inc., San Diego, CA; MSX2-AD) modified by the addition of two metabolic feeder hoods (Laboratory Products, Inc., Seaford, DE; 2110S) that prevent mice from caching powdered diets, and thus enable the tracking daily macronutrient intake (see below). Due to a lack of similar studies on this topic, we were unable to conduct an *a priori* power analysis to justify the number of donor/recipient mice per group. Instead, we decided on n = 10 per group based on the number of animals typically used in studies involving germ-free mice, the vast majority of which used 5-10 individuals per group^13^. One recipient mouse from the Herb-CONV group (V57) was excluded from our dataset due to aberrant behaviors that indicated possible injury during microbiome transplants. All recipient fecal samples were screened for 21 of the most common rodent pathogenic microorganisms using PCR tests conducted by a third-party diagnostic company (Charles River Research Animal Diagnostic Services, Wilmington, MA).

### Diet selection experiment

After colonization, conventionalized mice were acclimated for 7 days (to allow the gut microbiome to stabilize^55^), during which they were offered only sterile water and a low protein:carbohydrate ratio diet (LPC [0.27]; Table S1), as this diet is rather similar to standard mouse chow. After acclimation (day 0), mice were briefly removed from their cages for a 200 μL blood draw for metabolomics analysis (see details below). Mice were weighed (rounded to nearest hundredth) and returned to empty cages to facilitate the collection of fresh fecal samples for 16S rRNA microbial inventories and shotgun metagenomics (see details below). Conventionalized mice were then presented with a choice between two isocaloric diets (Table S1): (1) the LPC (0.27) diet offered during acclimation and (2) a diet with a higher P:C ratio (HPC [0.71]). The positions of these two diets were rotated daily to avoid learned preferences. Diets were designed by Teklad/Envigo (Indianapolis, IN), and were powdered prior sterilization to be visually indistinguishable from each other and to prevent food caching. Daily food consumption was calculated as the difference between the mass (rounded to nearest thousandth) of each diet presented (∼8 g) and the mass of each diet remaining after a 24-hour period. After tracking diet preferences for 11 consecutive days, animals were euthanized and dissected to investigate differences in the empty masses (rounded to nearest thousandth) of intestinal compartments. Conventionalized mice were maintained on a 12:12-h light:dark cycle, with 21°C ambient temperature and 40% humidity for the duration of the experiment. Animal experiments were conducted at the University of Pittsburgh Plum Borough Primate Facility under IACUC protocol 19074445.

### Metabolomics

Blood plasma was analyzed for primary metabolites (amino acids, hydroxyl acids, carbohydrates, sugar acids, sterols, aromatics, nucleosides, amines, and miscellaneous compounds) by the West Coast Metabolomics Center at the University of California – Davis, which performed all sample preparation, data acquisition, and data processing as previously described^57^. Briefly, metabolites were extracted using a mixture of acetonitrile:isopropanol:water (3:3:2, v/v/v) as well as 1:1 acetonitrile:water for removal of protein from serum. Dried metabolite extracts were resuspended in methoxyamine hydrochloride in pyridine for derivatization before being analyzed using gas chromatography-time-of-flight (GC-TOF) using a LECO Pegasus IV mass spectrometer equipped with automated liner exchange (ALEX; Gerstel corporation) and cold injection system (CIS; Gerstel corporation) for data acquisition. The CIS temperature was set at 50□°C to 250□°C final temperature at a rate of 12 °C s−1. Raw GC-TOF MS data were preprocessed with ChromaTOF (version 2.32) and apex masses were used to identify metabolites using the BinBase database. Values were reported as peak height for the quantification ion (*m/z* value) at the specific retention index, which is more precise than peak area for low abundant metabolites. All database entries that were positively detected in more than 10% of the samples of a study design class for unidentified metabolites were reported. Raw peak heights were vector normalized to reduce the impact of between-series drifts of instrument sensitivity, caused by machine maintenance status and tuning parameters.

### DNA extractions

DNA was extracted from donor cecal contents and day 0 conventionalized mouse feces using the Qiagen PowerFecal DNA Kit (Qiagen, Hilden, Germany; 12830) following the manufacturer’s instructions.

### 16S rRNA microbial inventories

Extracted DNA from conventionalized mice and donor cecum contents was amplified and sequenced by the Genome Research Core of the University of Illinois at Chicago as previously described^58^. Briefly, polymerase chain reaction (PCR) was used to amplify a portion of the bacterial 16S rRNA gene for Illumina sequencing using the Earth Microbiome Project primers 515F (GTGCCAGCMGCCGCGGTAA) and 806R (GGACTACNVGGGTWTCTAAT) targeting the V4 region of microbial small subunit ribosomal RNA gene^59^. Amplicon libraries were sequenced using a 2×251 paired-end run on an Illumina MiSeq. In addition to donor and recipient fecal samples, we sequenced five ‘blank’ extractions to control for the possibility of microbial contamination during the extraction procedure and microbial DNA present in commercial extraction kits^60^. A total of 1,398,994 raw Illumina sequencing reads (mean of 22,206 per sample (n = 63) ± 1111 SE) were paired and quality filtered via the DADA2 pipeline^61^ in QIIME2 (version 2020.4)^62^ using default parameters. Sequences that passed the quality filter were clustered into amplicon sequence variants (ASVs), which were identified using the SILVA reference database (release 138)^63^. Identified ASVs were filtered to exclude non-bacterial sequences (archaea, chloroplast, eukaryote, and mitochondria), reducing our total number of reads to 1,396,450 (mean of 22,166 per sample ± 1,112 SE) and 4,359 ASVs. We detected a total of 4,118 ASVs in donor and recipient fecal samples, 19 (0.46%) of which were also detected in blank extractions (total of 260 ASVs from 27,807 reads with mean of 5,561 per sample ± 1,419 SE). As recommended by McMurdie and Holmes (2014)^64^, we used un-rarefied ASV tables for comparisons of colonization efficiency (Bray-Curtis distances), alpha diversity (ASV richness and Faith’s phylogenetic diversity), and beta diversity (Bray-Curtis and unweighted/weighted UniFrac distances^65^).

### Shotgun metagenomics

Extracted DNA from conventionalized mice was sent to CoreBiome, Inc. (St. Paul, MN) for shotgun metagenomic analysis using BoosterShot™□. Briefly, sequencing libraries were prepared using a procedure adapted from the Illumina Nextera Library Prep Kit (Illumina, 20018705) and sequenced on an Illumina NovaSeq using single-end 1×100 reads with the Illumina NovaSeq SP reagent kit (Illumina, 20027464). A total of 122,190,150 raw sequence reads (mean of 4,213,453 per sample (n = 29) ± 151,158 SE) were filtered for low quality (Q-Score < 30) and length (< 50), trimmed of adapter sequences, and converted into a single fasta using SHI7 (version 0.99)^66^. Sequences were then trimmed to a maximum length of 100 bp and aligned using BURST (version 0.99.8)^67^ at 97% identity against CoreBiome’s Venti database consisting of all RefSeq bacterial genomes with additional manually curated strains as well as a bacterial KEGG^68^ annotated database created from dereplicating the bacterial genes within the Venti database. KEGG orthology counts were converted to relative abundance within a sample and collapsed into KEGG modules for statistical analysis.

### Statistics

Differences in macronutrient and total diet intake across treatment groups were tested using a multivariate analysis of variance (MANOVA) while controlling for the effects of body mass. A *post hoc* power analysis for MANOVA was conducted using G*Power^69^ (version 3.1) to confirm that statistical power was sufficiently greater than the widely-accepted minimum threshold of 0.80^70^. Microbial community structure (from 16S rRNA inventories) was visualized using principal coordinates analysis (PCoA) on ASV relative abundances, which were then assessed for differences (controlling for multiple comparisons using false discovery rate corrected P-values) across treatment groups using non-parametric permutational multivariate analysis of variance (PERMANOVA), analysis of similarity (ANOSIM), and permutational analysis of dispersion (PERMDISP) in QIIME2^62^. Microbiome function was visualized using PCoA on KEGG module relative abundances and analyzed for differences across treatment groups with PERMANOVA in QIIME2. Differences in the relative abundance of functional KEGG modules across conventionalized mice were tested using the non-parametric Krustal-Wallis test and linear discriminant analysis in LEfSe using the “one-against-all” strategy for multi-class analysis^71^. Identified plasma metabolites were filtered (based on mean intensity and IQR) and auto-scaled before using non-parametric median tests to identify metabolites that varied significantly across treatment groups and visualized using supervised partial least square discriminant analysis (PLS-DA) in MetaboAnalyst (version 4.0)^72^. Non-parametric Spearman rank correlations between plasma Trp availability, Trp KEGG modules, and macronutrient intake were conducted using non-parametric Spearman’s test (controlling for the effect of donor species) in the R package *ppcor* (version 1.1)^73^ and visualized using *corrplot* (version 0.85)^74^. Differences in empty cecum mass, empty colon mass, and colon length across treatment groups were tested using ANOVA with body mass as a covariate and corrected for multiple comparisons using Tukey’s HSD. Unless otherwise noted, all statistical tests were two-sided and conducted in JMP Pro version 14.1.0 (SAS Institute Inc., Cary, NC). For all statistical analyses, P-values ≤ 0.05 were defined as ‘significant’.

