## Supplementary figures and images for "The gut microbiome influences host diet selection behavior"

### Fig. S1

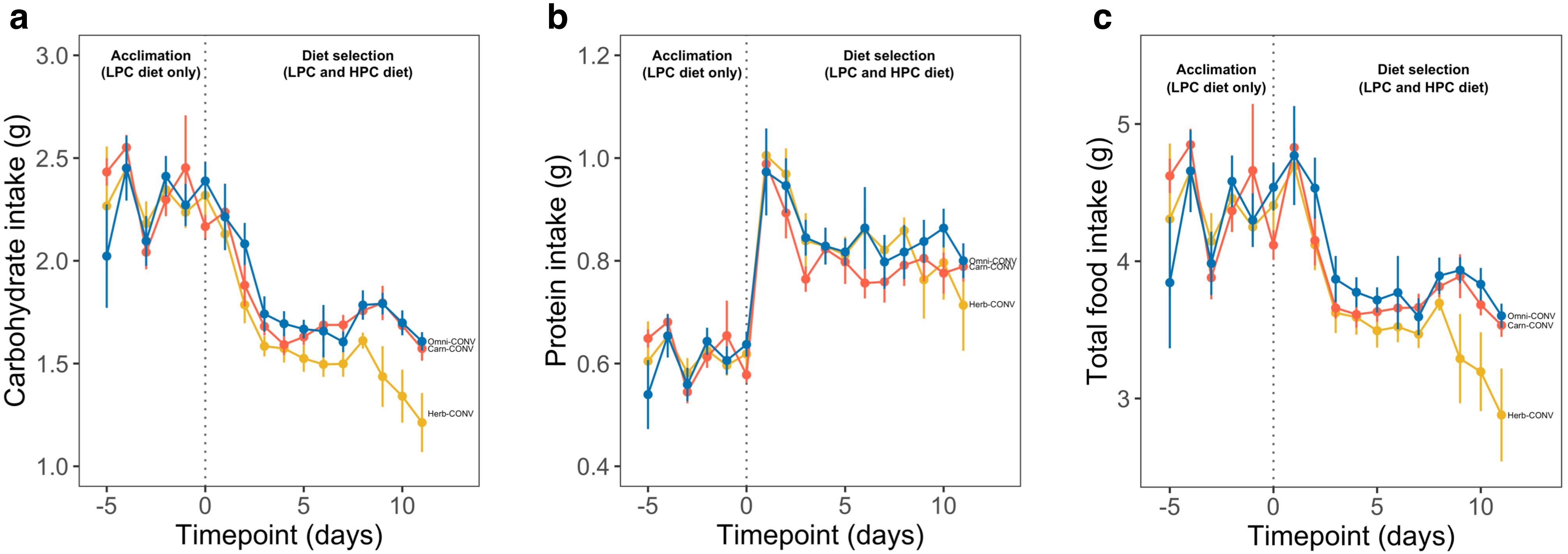

### Fig. S2

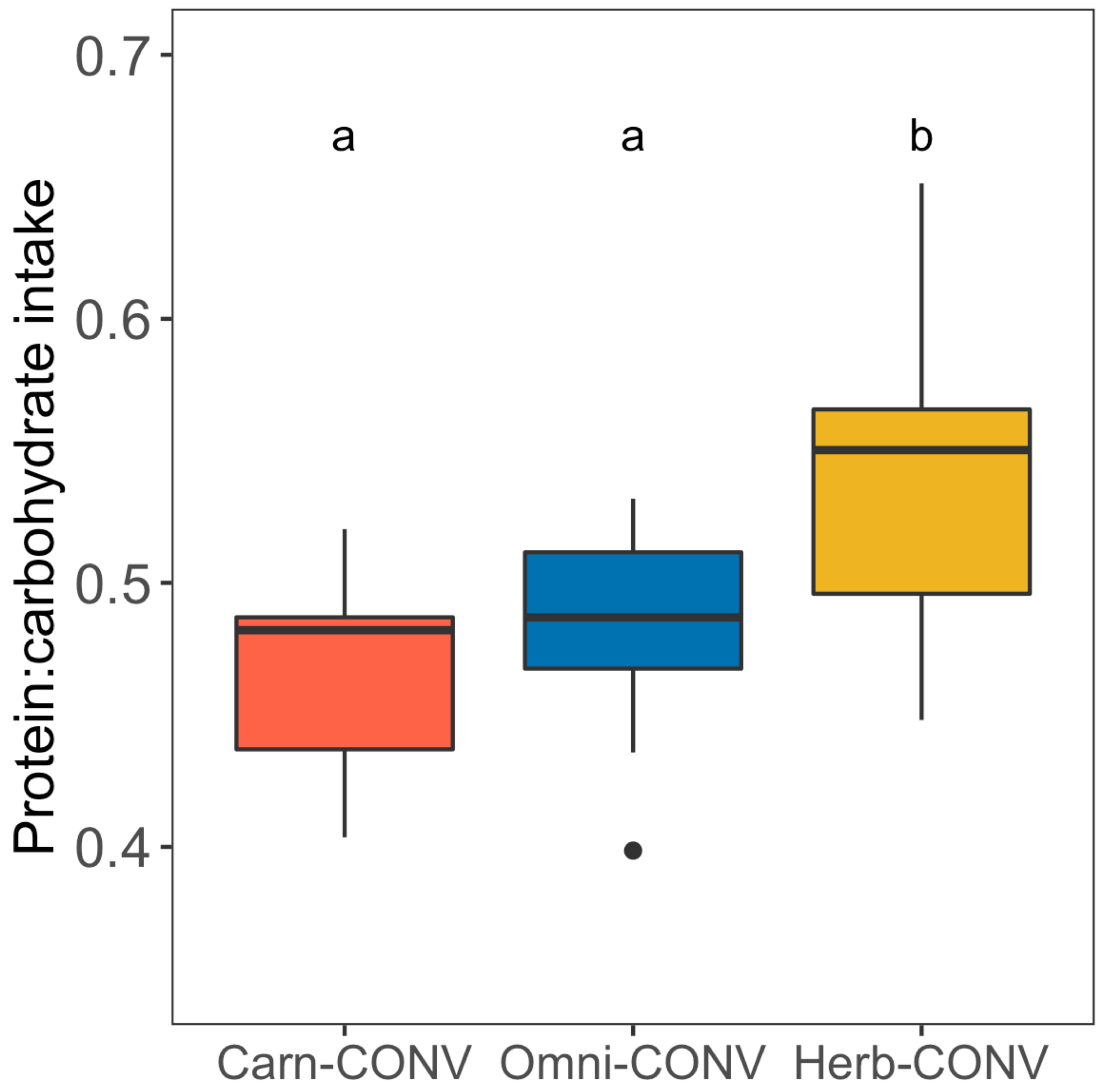

### Fig. S3

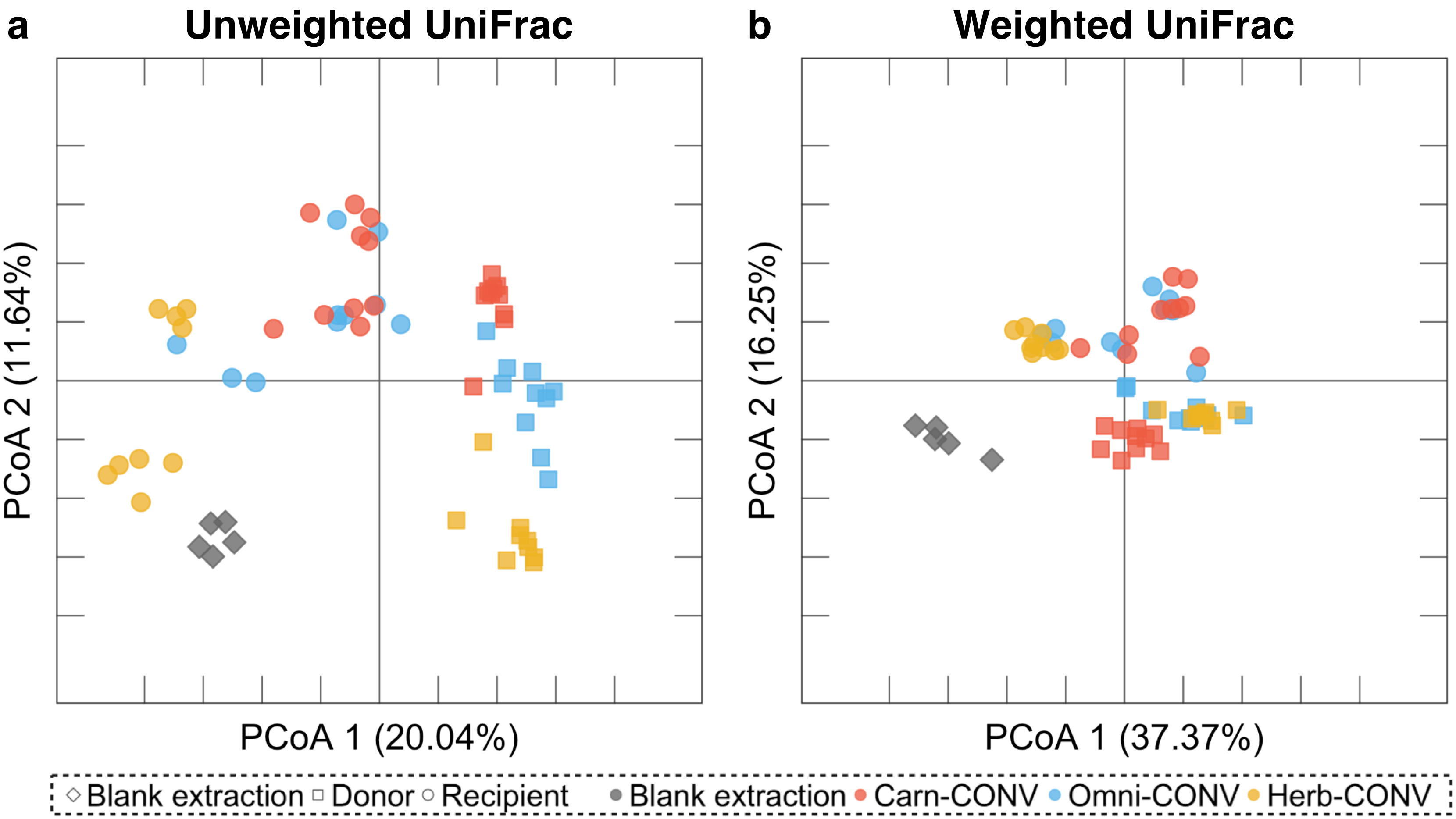

### Fig. S4

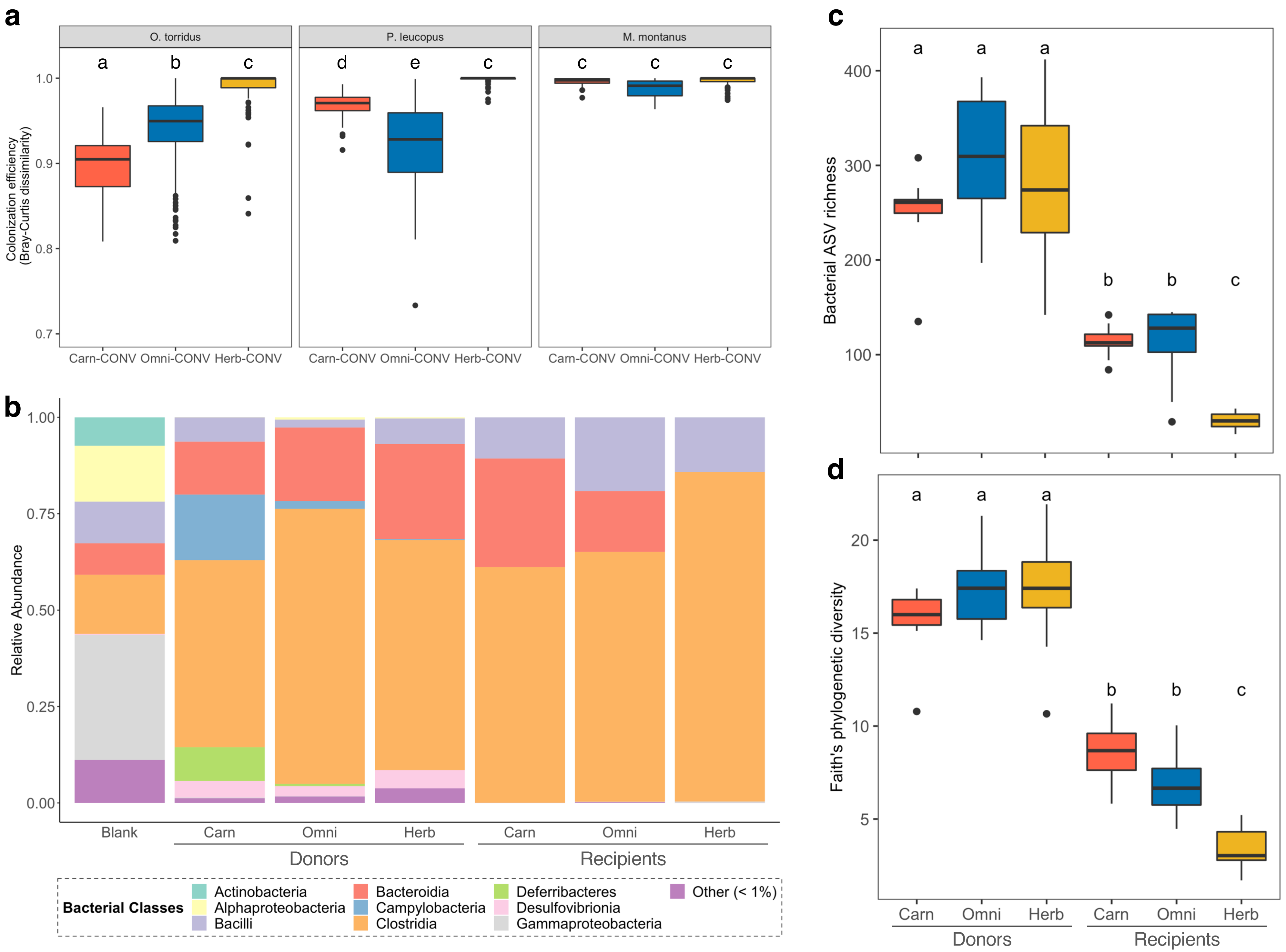
