## Supplementary material for "The gut microbiome influences host diet selection behavior": Table S1

| Diet composition | Low Protein:Carbohydrate diet (LPC) | High Protein:Carbohydrate diet (HPC) |
|---|---|---|
| Protein (g/Kg) | 140.39 | 277.44 |
| CHO (g/Kg) | 526.23 | 392.23 |
| Fat (g/Kg) | 50.28 | 49.75 |
| Fiber (g/Kg) | 32.90 | 40.65 |
| NDF (g/Kg) | 120.09 | 119.51 |
| Ca (g/Kg) | 9.99 | 10.01 |
| Cl (g/Kg) | 2.77 | 2.44 |
| K (g/Kg) | 8.23 | 7.72 |
| Mg (g/Kg) | 1.67 | 1.47 |
| Na (g/Kg) | 1.36 | 1.41 |
| P Avail (g/Kg) | 3.48 | 3.47 |
| P (g/Kg) | 5.64 | 5.11 |
| B-12 (mg/Kg) | 0.03 | 0.03 |
| B-6 (mg/Kg) | 21.65 | 20.47 |
| Biotin (mg/Kg) | 0.56 | 0.56 |
| Folic Acid (mg/Kg) | 2.36 | 2.35 |
| Niacin (mg/Kg) | 136.36 | 129.75 |
| Pantothenate (mg/Kg) | 68.42 | 67.71 |
| Riboflavin (mg/Kg) | 23.43 | 23.37 |
| Thiamin (mg/Kg) | 19.56 | 19.84 |
| Vit A (IU/Kg) | 19856.00 | 19888.00 |
| Vit D (IU/Kg) | 2204.50 | 2206.50 |
| Vit E (IU/Kg) | 143.92 | 138.69 |
| Vit K (mg/Kg) | 50.07 | 50.01 |
| Choline (mg/Kg) | 2074.46 | 2045.11 |
| Inositol (mg/Kg) | 1128.92 | 1372.32 |
| PABA (mg/Kg) | 110.13 | 110.13 |
| Vit C (mg/Kg) | 991.19 | 991.19 |
